## Supplemental Figures for "Parasitic nematode fatty acid- and retinol-binding proteins compromise host immunity by interfering with host lipid signaling pathways"

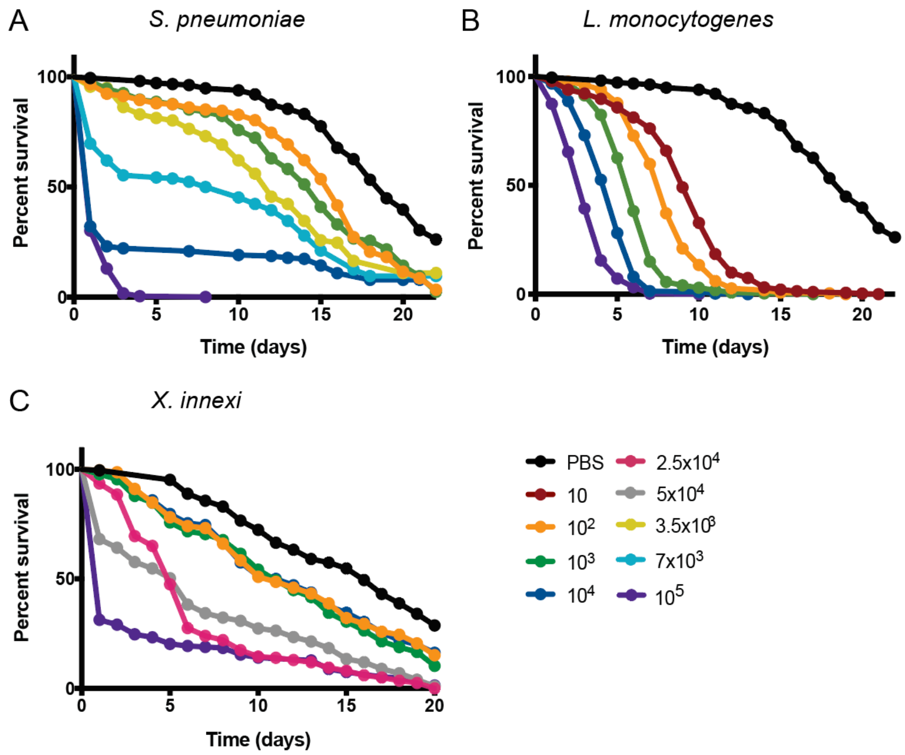


S-Fig 1: Dose response of various pathogens in wild-type OregonR flies was used to assess virulence patterns and to identify the $LD_{30}$, (dose that leads to 30% death of the population within the first 1-5 days depending on the pathogen) an optimal dose to measure variations in immunity. 5-7-day-old male flies were used for all injections with phosphate buffered saline (PBS) shown as the vehicle control. A) 100 to 100,000 CFUs of *Streptococcus pneumoniae* were injected, $LD_{30}$ identified as 7,000 cells. B) 10 to 100,000 cells of *Listeria monocytogenes* were injected, $LD_{30}$ identified as 1,000 cells. C) 100 to 100,000 cells of the insect pathogen *Xenorhabdus innexi* were injected, 25,000 and 50,000 cell doses were chosen for study. All raw data available in supplemental materials.

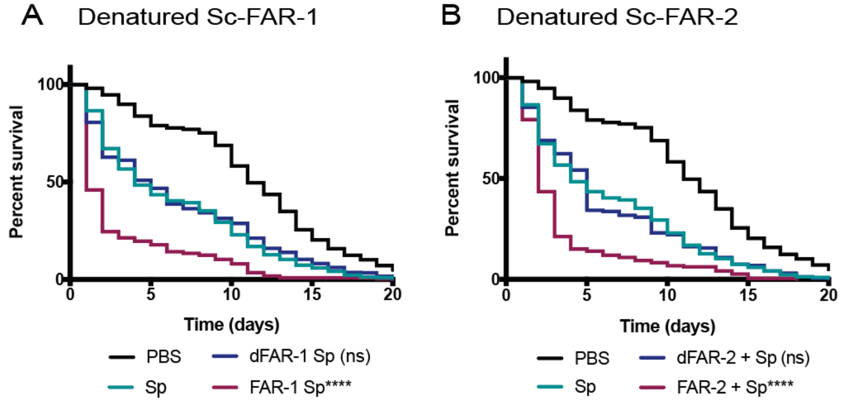


S-Fig 2: Recombinant Sc-FAR elicits a specific effect on the outcome of a bacterial infection. As a control to validate the effects of recombinant FAR proteins on immunity, denatured *S. carpocasae* FARs were co-injected with 7,000 CFUs of *S. pneumoniae*. A) Denatured Sc-FAR-1 (250ng) co-injected with *S.p.* shows no significant difference from *S.p.*-only injected flies. B) Denatured Sc-FAR-2 (250ng) co-injected with

*S.p.* shows no significant difference from *S.p.*-only injected flies. Statistics shown as Log-rank test. All raw data available in supplemental materials. Experiments not found to be significantly different from bacteria-only controls were marked ns.

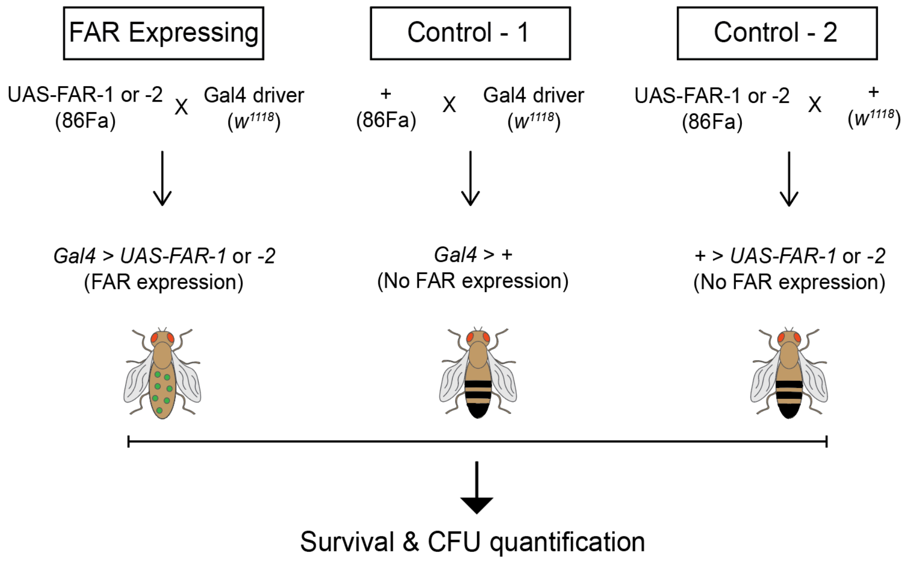


S-Fig 3: Overview of the UAS-Gal4 genetic crosses of FAR expressing and control flies. The sequence of FAR-1 or -2 was inserted into the 86Fa fly strain which displays a fluorescent red body and eye color and crossed with various Gal4 promoter strains. As a control, the 86Fa fly was crossed with the Gal4 driver flies as well as the FAR-1 or -2 transgenic flies were crossed with a *w*$^{1118}$ strain. The male flies yielding the appropriate genotypes were tested for their immunodeficiencies after inducting an immune response with a bacterial injection into the abdomen.

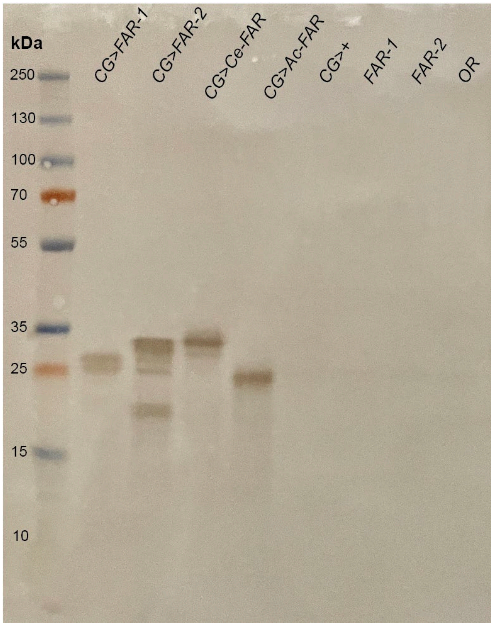


S-Fig 4: Western blot shows *in vivo* production of FAR proteins. FAR is expressed when driven with the Gal4 driver in the first 4 wells (rows 1-4). Control flies not expressed with UAS-Gal4 expression system do not show the presence of FAR proteins (rows 5-8).

A Natural Survival

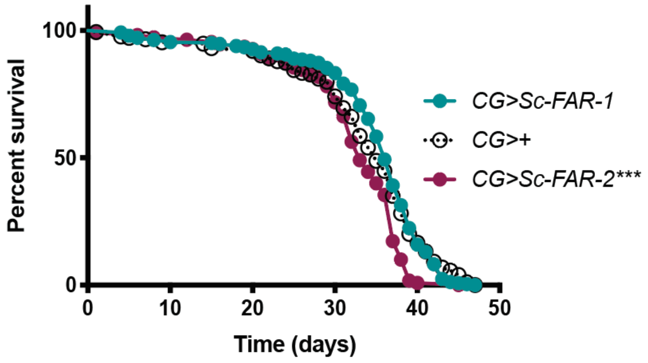


S-Fig 5: Lifespan of Sc-FAR transgenic flies expressed with the fat-body and hemocytes specific driver CG. Lifespan is not altered within the timeframe where immunity studies took place (day 0 to 20). Statistics shown as Log-rank test. All raw data available in supplemental materials.

A *$w^{1118}$ S. pneumoniae*

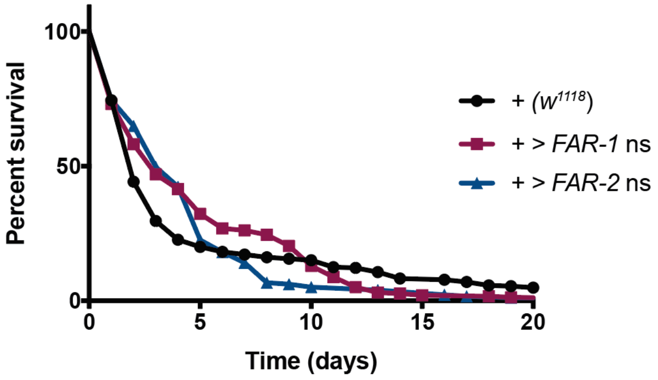


S-Fig 6: Genetic control of 86Fa transgenic strains. As a control to validate the specific effects on immunity after promoting the FAR transgenics, FAR-1 & -2 expressing flies were crossed with *$w^{1118}$* flies. There was no significant difference observed between all genotypes validating the specific effects of FAR. All raw data available in supplemental materials.

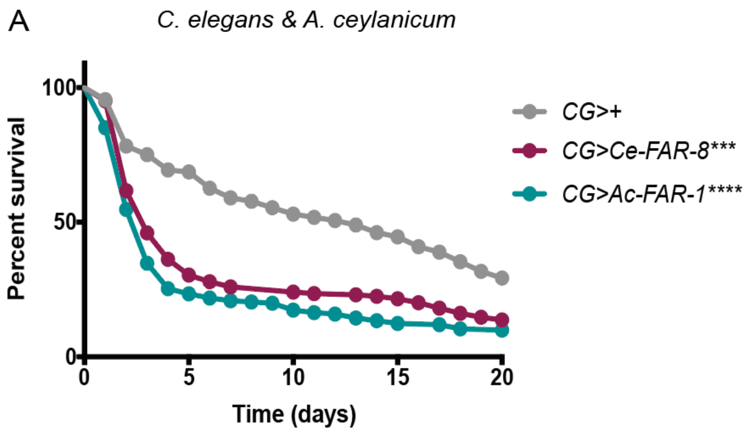


S-Fig 7: FAR proteins from other species of nematodes show similar negative effects on the outcome of a *S. pneumoniae* infection. Flies expressing FAR-8 from the free-living nematode *Caenorhabditis elegans* or a FAR from the mammalian hookworm *Ancylostoma ceylanicum* (maker-ANCCEYDFT_Contig87-pred_gff_fgenesh-gene-3.1) with the CG driver had a significant decrease in survival after injected with 7,000 cells of *S.p.* Statistics shown as Log-rank test. All raw data available in supplemental materials.

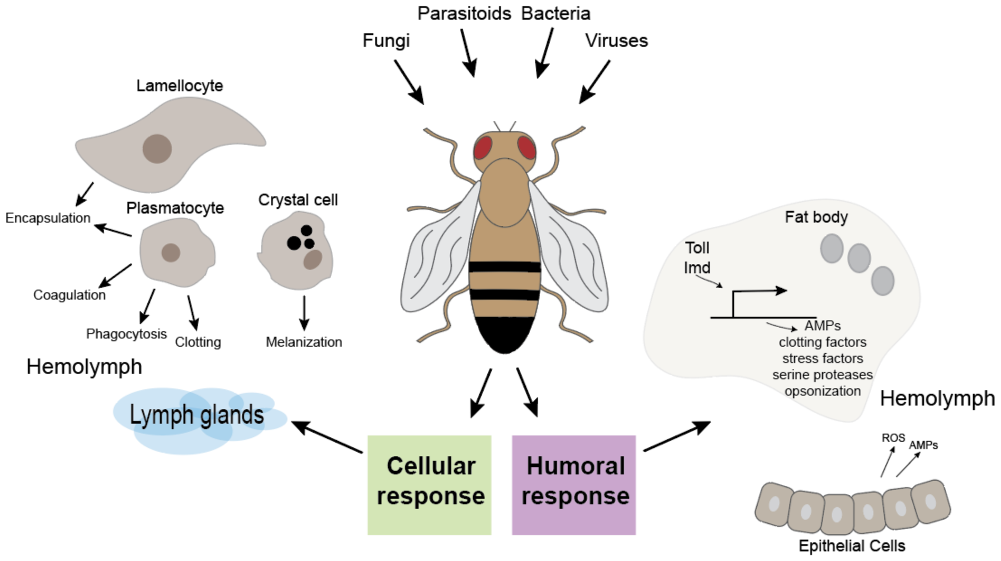


S-Fig 8: Simplified overview of Drosophila immunity. Detection of pathogens elicits an array of interconnected innate immune responses specifically divided into the humoral, or systemic, and the cellular response. Humoral immunity leads to the production of antimicrobial peptides (AMPs) downstream of either the Toll or Imd pathways. Cellular immunity is carried out by different types of hemocytes that surround and kill invading microbes. Adapted from (Lemaitre and Hoffmann, 2007; Vanha-Aho et al., 2016).

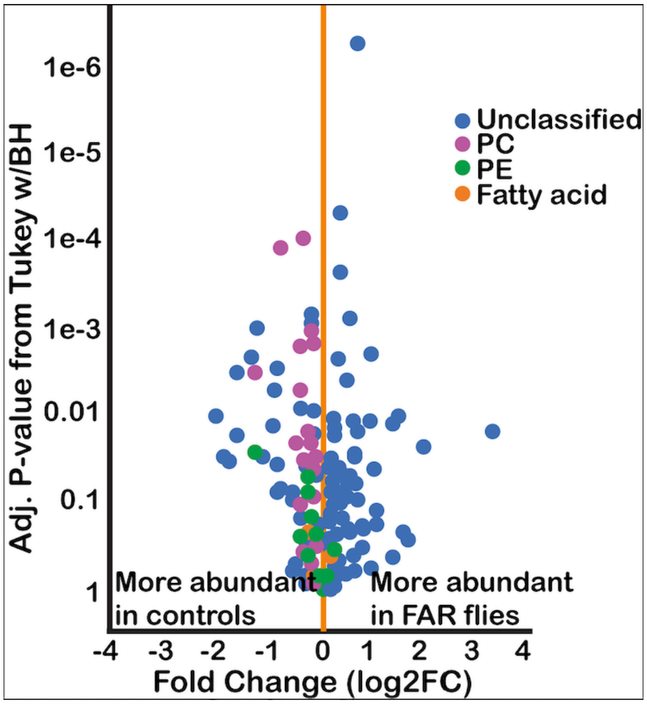


S-Fig 9: Volcano plot comparing relative abundance of metabolites in *CG>FAR-1* expressing and *CG>+* control flies. Phosphatidylcholines (PC), phosphatidylethanolamines (PE) and multiple fatty acids were found to be more abundant in control flies, meaning they are depleted in FAR expressing flies as well as potential binding partners, or upstream compounds of FARs binding partners. All raw data available in supplemental materials. LC-MS analysis depicted in this figure was performed at the UC Riverside Metabolomics Core Facility.

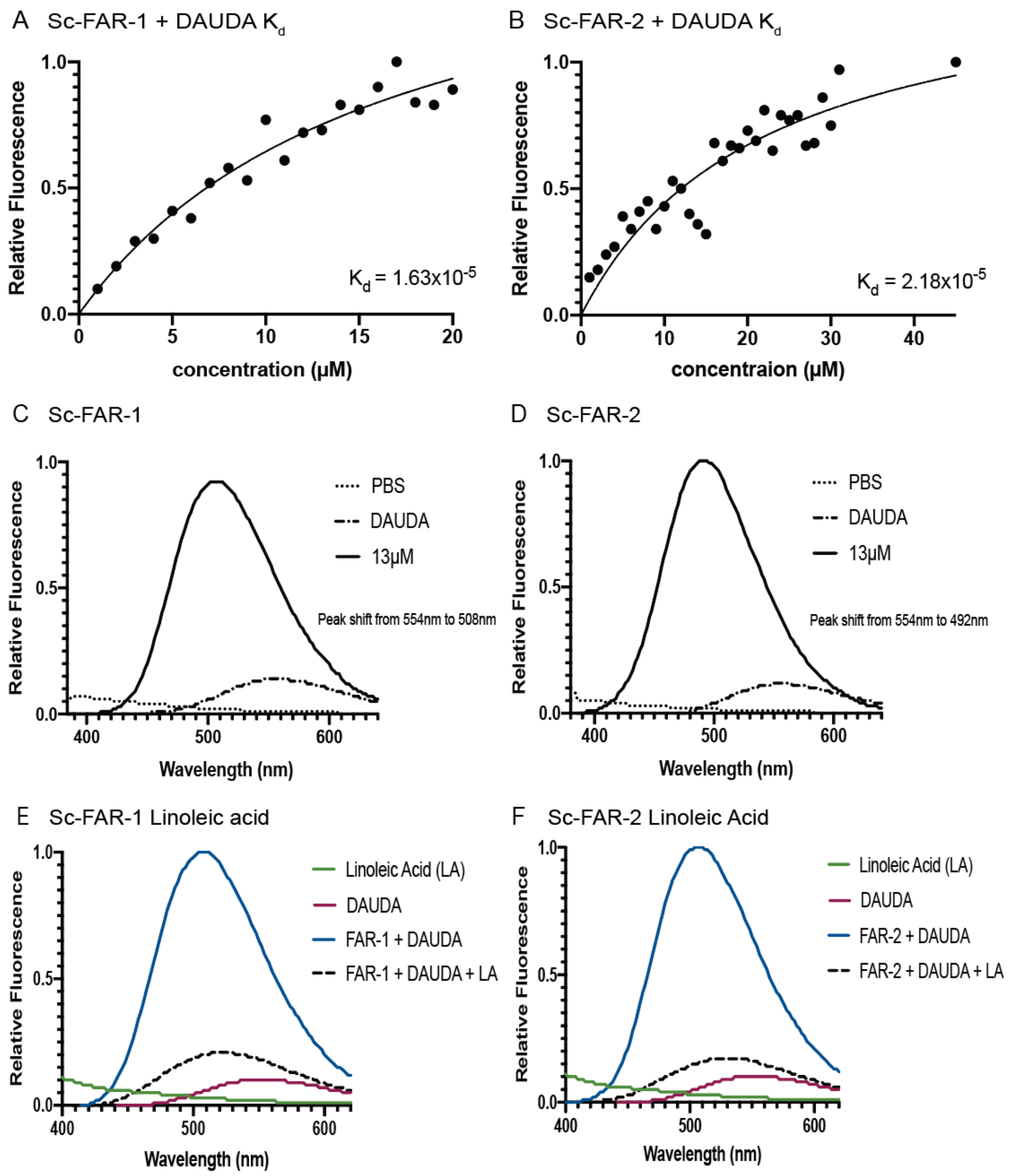


S-Fig 10: *In vitro* binding properties of Sc-FARs to fatty acids. A&B) Plots of increasing concentration of Sc-FAR binding to 1μM 11-(Dansylamino) undecanoic acid (DAUDA) in PBS reveal a K$_d$ in the 10 μM range. K$_d$ was estimated by using the best-fit in Graphpad Prism. C&D) When DAUDA is bound to FAR (13 μM) *in vitro*, a ~50nm blue shift is observed in the peak. E&F) Full curve of *in vitro* competition assays shows a decrease in fluorescence when linoleic acid (10 μM) binds to and therefore displaces DAUDA (1 μM) from the fatty-acid binding pocket of FAR. All raw data available in supplemental materials.

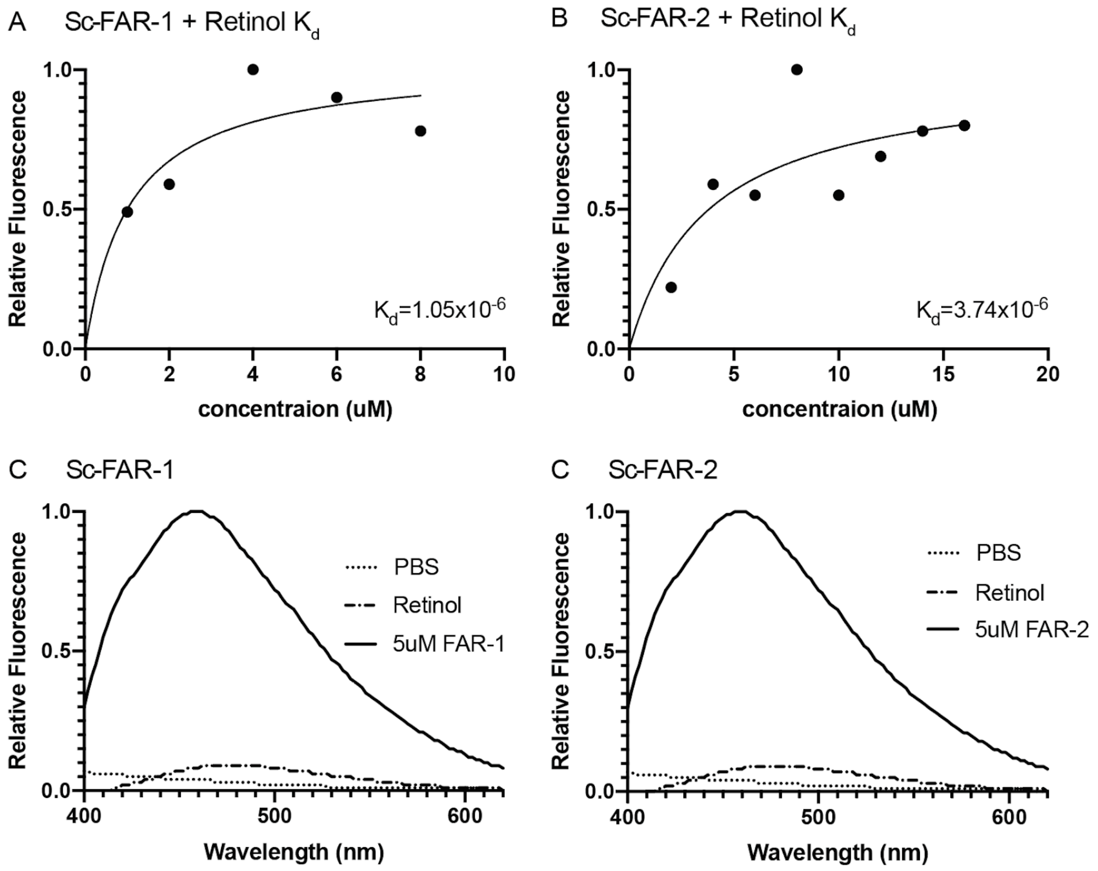


S-Fig 11: *In vitro* binding properties of Sc-FARs to retinol. A&B) Plots of increasing concentration of Sc-FAR binding to 40μM retinol in PBS reveal a $K_d$ in the 1 μM range. $K_d$ was estimated by using the best-fit in Graphpad Prism. C&D) When retinol is bound to FAR (5 μM) *in vitro*, the peak fluorescence is greatly increased. All raw data available in supplemental materials.

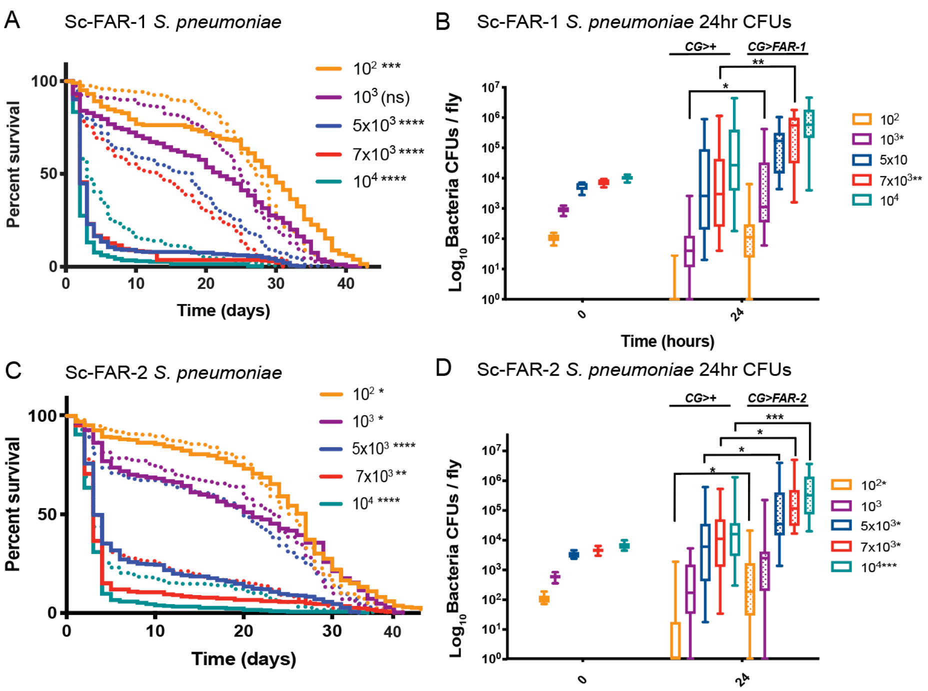


S-Fig 12: FARs' role in increased mortality and microbe load is not limited to the $LD_{30}$ dose. Sc-FAR-1 and -2 were expressed with the CG driver and injected with 100 to 100,000 CFUs of *S.p.* A&C) Survival was measured daily showing that Sc-FARs significantly increase mortality rate in most doses. B&D) In the presence of FAR, various doses of *S.p.* also cause an increase in microbe growth 24 hours p.i. Dotted lines show *CG>+* and solid lines show *CG>FAR-1* or *-2.* Statistics shown as Log-rank tests for survival curves and unpaired t-tests for microbe growth. All raw data available in supplemental materials.
